# We are not ready yet: limitations of transfer learning for Disease Named Entity Recognition

**DOI:** 10.1101/2021.07.11.451939

**Authors:** Lisa Langnickel, Juliane Fluck

## Abstract

Intense research has been done in the area of biomedical natural language processing. Since the breakthrough of transfer learning-based methods, BERT models are used in a variety of biomedical and clinical applications. For the available data sets, these models show excellent results – partly exceeding the inter-annotator agreements. However, biomedical named entity recognition applied on COVID-19 preprints shows a performance drop compared to the results on available test data. The question arises how well trained models are able to predict on completely new data, i.e. to generalize. Based on the example of disease named entity recognition, we investigate the robustness of different machine learning-based methods – thereof transfer learning – and show that current state-of-the-art methods work well for a given training and the corresponding test set but experience a significant lack of generalization when applying to new data. We therefore argue that there is a need for larger annotated data sets for training and testing.

## Introduction

The amount of freely available, electronic data increased enormously in the biomedical field. Automatic information extraction methods have become indispensable and intense research has been done in the past. Whereas most text mining tasks were achieved with the help of rule-based systems in the beginning, mainly machine learning methods are used nowadays. The latter are strongly dependent on large amounts of curated data. However, manual curation is a complex and time consuming task, at least in the biomedical field, that needs to be done by domain experts. Hence, the availability of such high-quality data sets is strongly limited. In the area of biomedical named entity recognition (NER), most data sets have been released for shared tasks and challenges open for the community. To name two examples, the national NLP clinical challenges (n2c2), formerly known as *i2b2 NLP Shared Tasks*, provide curated clinical data to researchers (1); the organization *Critical Assessment of Information Extraction systems in Biology (BioCreAtivE)* organizes challenges for biological natural language processing (NLP) tasks and therefore also releases annotated data. In terms of disease entity recognition, to the best of our knowledge, two publicly available literature data sets exist that are commonly used: the NCBI Disease corpus (2) and the BC5CDR Disease corpus (3). Both of the mentioned disease data sets follow the same annotation guidelines which are necessary to ensure consistency in annotations. These guidelines have been published together with the NCBI Disease corpus^1^ and are also used for the more recent one (BC5CDR)^2^.

The machine learning-based approaches applied to NLP have changed over time. First, methods like *support vector machines*, *hidden markov models* or *conditional random fields*, which all belong to the class of supervised algorithms, where often superior compared to rule based approaches. For those techniques, so-called features are needed to describe the input data. Examples of used features include general linguistic features (e.g. part-of-speech tags, stems), orthographic features (e.g. punctuation character, capitalized word) or dictionary look-up features. Later, so-called word embeddings replaced this feature engineering process. Word embeddings are “representations of words in n-dimensional space, typically learned over large collections of unlabeled data through an unsupervised process” with the help of neural networks (4). These vectors are usually pre-trained with the objective to build a general language model, i.e. to predict the next word in a sequence. This principle can be understood as providing the neural network with prior knowledge about the nature of words and sentences – i.e. their semantics and syntax. The aforementioned methods are all feature-based approaches: pre-trained representations (word embeddings) are included as features for a task-specific architecture (5). More recently, so-called fine-tuning approaches have gained interest, which exploit a mechanism known as transfer learning. A trained model is used as starting point to be trained on a new task. In case of NLP, the model is pre-trained on a general language understanding task and then fine-tuned on a specific NLP task like NER or relation extraction (RE). With this shift in text mining methodologies, the complexity of the workflow is drastically reduced compared to rule- and feature-based approaches. Rule-based approaches require several pre-processing steps as for instance part-of-speech (POS) tagging, tokenization and sentence detection. Feature-based approaches rely on at least two different architectures, i.e. the creation of features and their inclusion into a (different) model.

In contrast, fine-tuning based approaches only define one network architecture that is applicable to several different downstream tasks. The most popular network architecture is the bidirectional encoder representation transformer (BERT) (5) that has been adapted to the biomedical area, called BioBERT, and shows state-of-the-art results for several different NLP tasks, thereof disease NER (6).

Based on the needs during the current COVID-19 pandemic, we set up the text mining-based semantic search engine preVIEW that automatically indexes preprints from several different sources (7). To recognize several entity classes (thereof diseases), we integrated publicly available ML-based models which show promising results of F1-scores above 85% for disease name recognition. Unfortunately, we realized a significant drop in performance when evaluated on a newly annotated COVID-19 preprint data set. However, with the implementation of an additional post-processing step, we could achieve good results for this specific corpus. These findings encouraged us to examine this performance reduction phenomenon in more detail – based on known data heavily used by the community: To the best of our knowledge, all recently developed systems for the recognition of diseases are trained and evaluated on either one of the above mentioned data sets, on both of them separately or on the combination of these data sets. The question arises whether the models trained on these data sets are robust and applicable to real world applications. In the current work, we investigate different algorithms and compare performance of the algorithms trained on data set *A* and tested on the data set *B*. This will be termed as *cross evaluation* in the following. We also investigate the similarities and differences of the two data sets and in addition compare them to a random PubMed data set in order to analyze the characteristics/bias of the different data sets.

## Materials and Methods

In the following, we first describe the two used data sets: the NCBI disease corpus and the BC5CDR disease corpus. Afterwards, we describe the used algorithms in more detail.

### Data sets

The two different data sets both consist of PubMed abstracts with manually curated disease annotations. The NCBI disease corpus with detailed annotation guidelines was released first. For the generation of the BC5CDR corpus, the previously published NCBI disease guidelines were re-used. The authors stated that “whenever possible, we will follow closely the guidelines of constructing NCBI disease corpus for annotating disease mentions” (8).

The NCBI Disease corpus was released by the National Center for Biotechnology Information (NCBI) and is “fully annotated at the mention and concept level to serve as a research resource for the biomedical natural language processing community” (2). It contains 739 PubMed abstracts with a total of 6,892 disease mentions, annotated by a total of 14 annotators. Two annotators were given the same data so that a double-annotation could be performed. The inter-annotator agreement was determined by means of the F1-score (see Evaluation metrics) for each pair of annotators. The average F1-score amounts to 88% (2).

The BioCreative V Chemical Disease Relation (BC5CDR) was released by the organization BioCreAtIvE. The BC5CDR corpus contains 1,500 abstracts including disease and chemical annotations at mention level as well as their interactions (relations). In total, the data set contains 12,848 disease mentions (3). For the present work, only the corpus containing disease mentions is used. Here, the inter-annotator agreement has been determined by means of the Jaccard distance. The Jaccard index divides the overlap of both sets (annotations) by the number in either set (9). To determine the Jaccard distance, the index needs to be subtracted from one. The inter-annotator agreement amounts to 87.49% (3).

The NCBI training data set consists of 593 and the BC5CDR training data set consists of 500 abstracts. In terms of unique mentions and concepts, they are also very similar. Whereas the NCBI training set contains 632 unique concepts, 649 can be found in the BC5CDR training set. In the test sets, huge differences can be found concerning the amount. The NCBI Disease test set only consists of 100 abstracts, the BC5CDR test set, however, consists of 500 abstracts as well. Therefore, the latter contains significantly more unique mentions and concepts. A detailed overview can be seen in Table 1.

**Table 1.** Statistics of used disease entity recognition data sets

|  | Data set | NCBI | BC5CDR |
| --- | --- | --- | --- |
| Amount of abstracts | Training | 593 | 500 |
| Unique mentions |  | 1614 | 1445 |
| Unique concepts |  | 632 | 649 |
| Amount of abstracts | Development | 100 | 500 |
| Unique mentions |  | 343 | 1343 |
| Unique concepts |  | 170 | 589 |
| Amount of abstracts | Test | 100 | 500 |
| Unique mentions |  | 407 | 1432 |
| Unique concepts |  | 192 | 640 |

In our work, we analyze and compare these data sets on different levels: on mention level, on concept level and based on the whole corpus. For the latter, we apply the tool *scattertext* to visualize the linguistic variations (10). In addition, we use the randomly generated PubMed corpus to perform a linguistic variation analysis between the annotated corpora and PubMed. This corpus was generated by randomly choosing 500 abstracts from all PubMed abstracts with a publication date between 1990 and 2021 (a total of 23,631,092 articles).

### NER Algorithms

We investigated four different publicly available algorithms for disease named entity recognition in this work, that will be described in the following. Whereas we trained BioBERT in this study, we applied the other algorithms “as is”. We provide an overview about the sources in the Availability Section.

BioBERT (6) is based on BERT (5) which, in turn is based on the transformer architecture that was developed by Vaswani *et al.* (11). The transformer architecture is based on an encoder-decoder structure trained on translation tasks. For building a bidirectional language model, Devlin *et al.* adapted only the encoder structure from the original transformer. In contrast to previously described methods to build a language model, BERT uses a different learning strategy. Instead of predicting each next (or previous token) only 15% of the words, which are chosen randomly, are masked and then predicted. The second task is a binary classification task predicting whether or not a sentence follows another given sentence. As pre-training corpus, two large corpora are combined, namely the BooksCorpus consisting of 800M words and the English Wikipedia corpus with 2,500 words. The BooksCorpus is a collection of 11,038 electronic books (12). BioBERT models are based on the pre-trained BERT model but are additionally trained on domain-specific corpora, i.e. PubMed and PMC articles. As pre-trained model, we used *BioBERT-Base v1.0 (+ PubMed 200K + PMC 270K)* published by Lee *et al.* (6). For fine-tuning, we used the library *Transformers* (13) and pytorch. We trained three different models, one on the NCBI Disease training data set, one on the BC5CDR training data set and one on the combination of them. For the last setting, the batches were shuffled randomly to avoid a higher influence of one data set over the other. The training parameters, investigated via cross-validation, can be seen in Table 2.

**Table 2.**
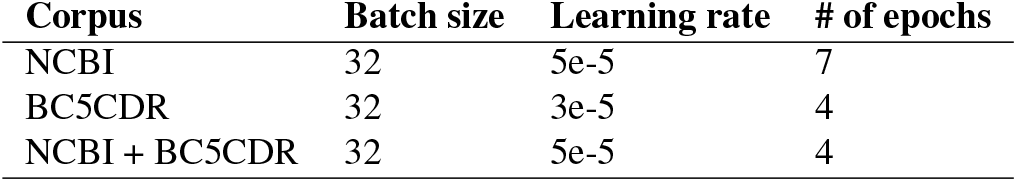
BioBert parameters

| Corpus | Batch size | Learning rate | # of epochs |
| --- | --- | --- | --- |
| NCBI | 32 | 5e-5 | 7 |
| BC5CDR | 32 | 3e-5 | 4 |
| NCBI + BC5CDR | 32 | 5e-5 | 4 |

scipaCy is based on the python library spaCy (14) that includes tools for text processing in several different languages. The text processing steps include for example sentence detection, tokenization, POS tagging or NER. Therefore, a convolutional neural network (CNN) is used. scispaCy is trained on top of spacy for POS tagging, dependency parsing and NER using biomedical training data. The authors provide a model trained on the BC5CDR corpus to recognize diseases and chemicals. We used this model and filtered out the chemical annotations.

DNorm is a disease recognition and normalization tool (15). It is a serial algorithm which uses first the entity recognition tool BANNER (16) based on conditional random fields (CRFs) which is followed by an abbreviation detection tool and a normalizer. Normalization is learned following a pairwise learning to rank approach. The authors provide different models optimized on either NCBI Disease corpus or BC5CDR Disease corpus. We apply both types of models and evaluate them on both disease data sets, respectively.

TaggerOne is a joint named entity recognition and normalization model consisting of “a semi-Markov structured linear classifier, with a rich feature approach for NER and supervised semantic indexing for normalization” (17). The authors provide three different models, trained on the NCBI Disease corpus, the BC5CDR Disease corpus and both of them simultaneously.

An overview about all available/developed models can be seen in Table 3.

**Table 3.** Overview of used disease entity recognition models

| Training set | Algorithm |  |  |  |
| --- | --- | --- | --- | --- |
|  | BioBERT | scispaCy | DNorm | TaggerOne |
| NCBI | ✓ |  | ✓ | ✓ |
| BC5CDR | ✓ | ✓ |  | ✓ |
| NCBI+BC5CDR | ✓ |  |  | ✓ |

### Evaluation Metrics

We determine precision, recall and F1-score to evaluate the models. The equations are given below, where FP stands for false positive, FN for false negative and TP for true positive. To ensure consistency, we use a publicly available evaluation script (CoNLLEval script) that has been released by the Conference on Computational Natural Language Learning (CoNLL) together with a shared task. The script is available under https://github.com/sighsmile/conlleval. This requires the input data to be in the “BIO”-format where each token is labeled as *B* for beginning, *I* for inside or *O* for outside.

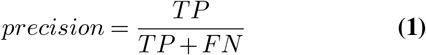

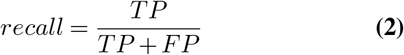

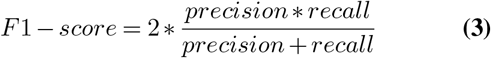

## Results

This section is subdivided into three different parts. First, we describe the results of the corpora comparison analyses. Afterwards, the results of the cross evaluation are described and finally we present the results of the combined learning approach.

### Semantic and linguistic comparison of data sets

We first determined the overlap of both mentions and concepts between the training and the corresponding test set. The overlap between NCBI Disease training and its test set reaches 70% on concept level, compared to an overlap of 60% between BC5CDR training and test set. Second, we determined the “cross-similarity”, i.e. the similarity of the training set of the NCBI disease corpus and the test set of the BC5CDR corpus and vice versa. The overlap between NCBI Disease training set and BC5CDR test set only reaches 32% on the concept level and for the opposite case a value of 24% is reached. An overview of all results, also on the mention level, is given in Figure 1. On the mention level, the overlap is lower within a corpus but we can also observe a drastic drop of cross-similarity.

In a next step, we compared the linguistic variability of the different corpora. In Figure 2a, we compared the BC5CDR training corpus to its corresponding test set. It shows a positive, linear relationship, indicating that the same words (or words with similar meaning) occur with similar frequency. In contrast, we do not see a relationship between the BC5CDR training set and the NCBI disease training set as the points are scattered throughout the whole plot (see Figure 2b). This means, that terms that occur often in the BC5CDR training set occur rarely in the NCBI training set and vice versa. Finally, we compared both, the NCBI and the BC5CDR corpus, to the random PubMed corpus and received similar results (see Figures 2c and 2d): in both cases also no linear trend can be seen but a widely distributed scatterplot.

**Fig. 1.**
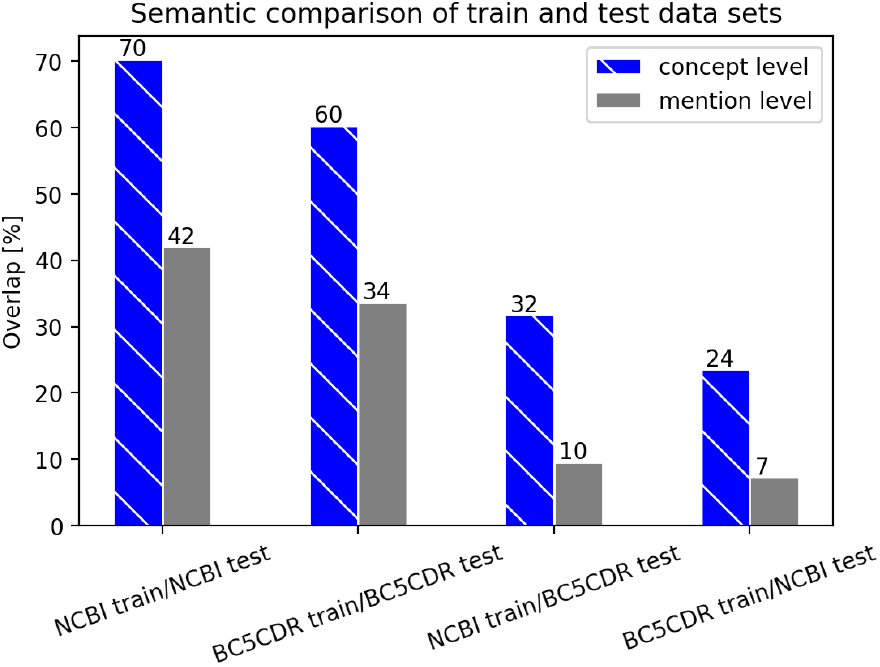
Comparison of training and test data set for the NCBI Disease and the BC5CDR corpus. The data sets are compared based on the overlap of mentions as well as concepts.

**Fig. 2.**
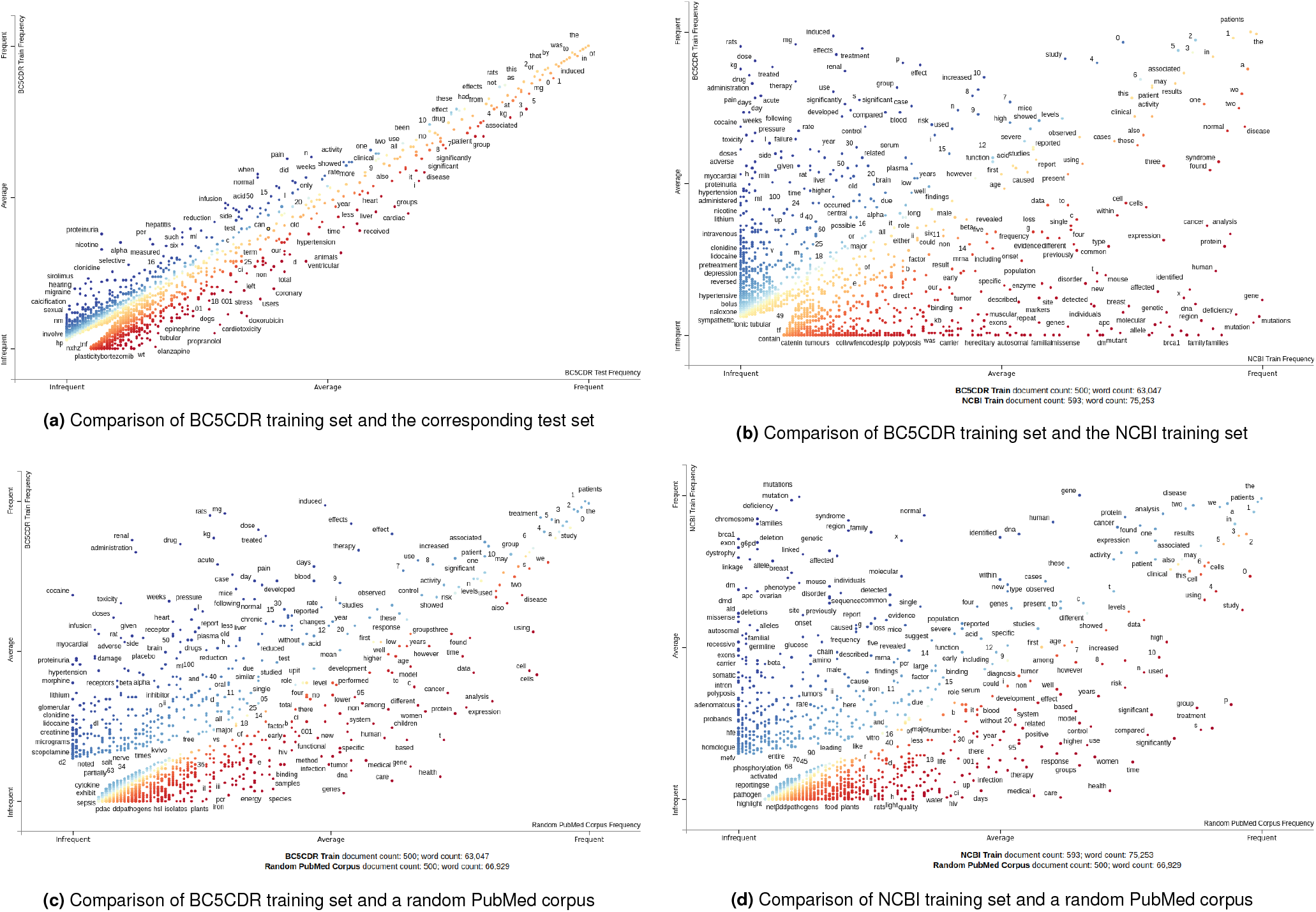
Comparison of the data sets with *scattertext*. On each axis, the frequency of a term is shown for the given documents. In Figure 2a, the BC5CDR training set is compared to its given test set whereas in Figure 2b, the BC5CDR training set is compared to the NCBI training set. In Figures 2c and 2d, the BC5CDR training set and the NCBI training set are compared against a randomly chosen PubMed corpus of similar size.

### Cross Evaluation of NER models

All mentioned machine learning methods are evaluated on both available test sets (NCBI and BC5CDR Disease). As can be seen in Figure 3, the cross evaluation results in a significant drop for all used models. Whereas the BioBERT model trained on the NCBI training corpus achieves an F1-score of about 87% on the corresponding test set, it drops to 68% for the BC5CDR Disease test set. Similarly, the BioBERT model trained on the BC5CDR training set reaches an F1-score of 83% on the corresponding test set, the cross-evaluation, however, results in an F1-score of 69%. The highest difference is determined for the TaggerOne model trained on the NCBI training set. Whereas an F1-score of 83% for the corresponding test set is achieved, only 52% are reached for the BC5CDR test set. Vice versa, for the TaggerOne model trained on the BC5CDR corpus, we realize a 20% drop for the cross-evaluation. For trained DNorm and scispaCy models, the same trend has been determined.

**Fig. 3.**
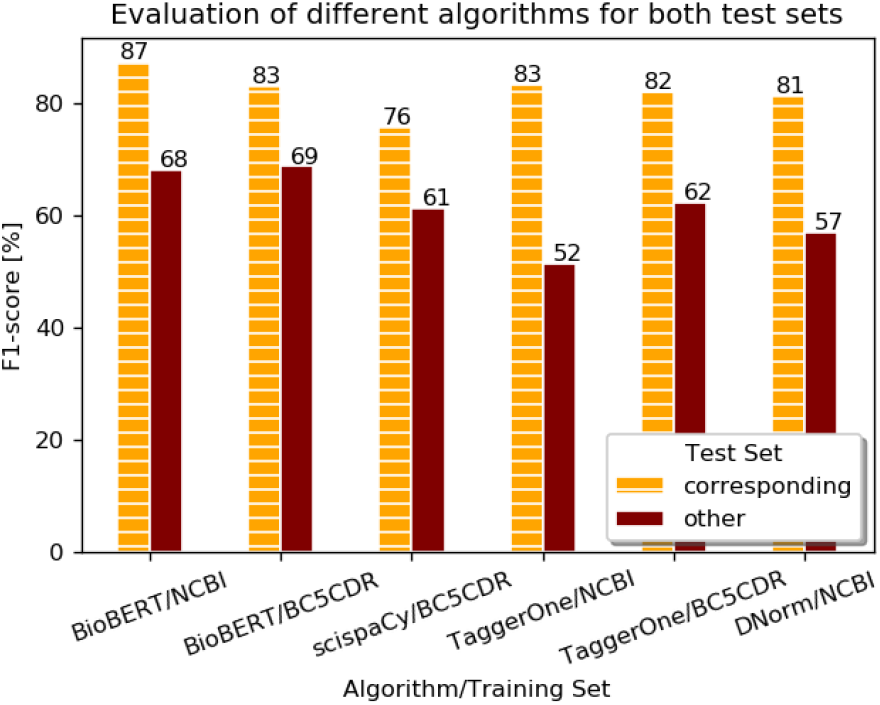
NER results for all tested ML algorithms. The F1-score is shown for the test set that belongs to the training set (corresponding test set) and to the test set of the respective other data set.

### Learning on combined data set

Finally, we trained a BioBERT model on both disease data sets simultaneously and also evaluated this on both test data sets. Also for TaggerOne such a combined model is provided that we evaluated. As can be seen in Table 4, this results are similarly high for both test data sets. For BioBERT, the result on the NCBI disease test set is only 0.07% worse than the model only trained on NCBI; the result on the BC5CDR disease test set is even the same (see Fig.3).

**Table 4.**
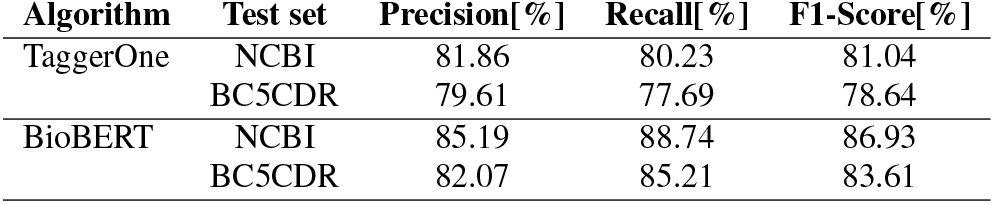
Evaluation of models trained on combined data set

## Discussion

In order to find relevant information in literature and hence to generate new knowledge, text mining methods have become indispensable because of the ever growing amount of electronic data. Therefore, a lot of research has been done in the area of bioNLP and current state-of-the-art algorithms show promising results on the available data sets. BERT is on everyone’s lips and used in a variety of biomedical and clinical applications (6, 18–20).

Because we integrated NER models into a semantic search engine and realized a drop in performance when evaluating an algorithm on a new data set, we started to question the robustness of current state-of-the-art methods. Therefore, in this work, we investigated the robustness of different machine learning-based algorithms on the task of disease named entity recognition. We chose this example because two different manually curated data sets are publicly available that basically follow the same annotation guidelines. Under the assumption that a machine or deep learning-based algorithm is able to generalize and to predict on new data, we evaluated a model trained on one of the available corpora on the test set of the other corpus. Our analysis shows that none of the four tested algorithms performs nearly as good on cross evaluation as on the corresponding test set. We experience a significant drop in performance – on average 20.5% in terms of F1-score. To our mind, this can have the following two reasons: (1) the models can be overfitted towards the training data sets or (2) one such available corpus is simply not enough to learn this kind of complex biomedical NLP task. As we showed in our scatterplots, the content of the two used data sets strongly differ in content and wording (see Fig.2). The model trained on the combination of the data sets reaches nearly the same results as each model trained on only one data set. We conclude that the BERT model is able to predict well on more variable test data if the training data set covers a similar variance. Therefore, the question arises when the model would be “ready” for real world applications – i.e. when we would have enough representative data. The model needs to be further tested on manually curated data that again covers a different area. However, such experiments are hampered by a lack of high-quality labeled data. Therefore, we foresee to set up a crowd sourcing-based approach in the near future and want to test the capabilities of transfer learning-based approaches for an active learning setup. However, sequential fine-tuning of BioBERT models (i.e. re-training) experiences a mechanism known as *catastrophic forgetting* – the model forgets previously gathered knowledge and is biased towards the last data set (21). Recently, so-called Adapter modules have been proposed that can be used for sequential learning of different tasks (22, 23). However, it remains open how such methods perform on exactly the same task (i.e. disease NER in our case).

## Conclusions

Even though current transfer learning-based state-of-the-art methods for bioNLP show excellent results on the given training and corresponding test data, our analysis showed that those models are – against our expectations – not yet ready for real world applications because of a lack of generalization capabilities. Named entity recognition in the biomedical domain is much more complex than solving tasks on general domain knowledge, such as the recognition of persons or organizations. Moreover, a continual learning process is of great importance as the science progresses not only continuously but also rapidly. Therefore, in our future work, we foresee both the manual annotation of further data sets and the investigation of continual learning capabilities on this task in order to be able to solve real world cases.

## Availability

- Scattertext: https://github.com/JasonKessler/scattertext
- BioBERT pre-trained models: https://github.com/naver/biobert-pretrained
- Transformers library to fine-tune BioBERT: https://github.com/naver/biobert-pretrained
- scispaCy library: https://allenai.github.io/scispacy/
- DNorm: https://www.ncbi.nlm.nih.gov/research/bionlp/Tools/dnorm/
- TaggerOne: https://www.ncbi.nlm.nih.gov/research/bionlp/tools/taggerone/
- Evaluation script: https://github.com/sighsmile/conlleval

## Footnotes

1 https://www.ncbi.nlm.nih.gov/CBBresearch/Dogan/DISEASE/Guidelines.html

2 https://biocreative.bioinformatics.udel.edu/media/store/files/2015/bc5_CDR_data_guidelines.pdf

